# DETECTION OF CELL ASSEMBLIES IN HIGH-DENSITY EXTRACELLULAR ELECTROPHYSIOLOGICAL RECORDINGS

**DOI:** 10.1101/2024.01.26.577338

**Authors:** Gabriel Makdah, Sidney I. Wiener, Marco N. Pompili

**Author notes:** Correspondence should be addressed to MNP.

## Abstract

**Cell assemblies, i.e., concurrently active groups of neurons, likely underlie neural processing for higher brain functions. Recent technological progress has enabled large-scale recording of neuronal activity, permitting the exploration and analysis of cell assembly dynamics. This review aims to provide both conceptual insights and practical knowledge pertaining to principal methodologies used for detecting cell assemblies in the last fifteen years. The goal is to assist readers in selecting and comparing various protocols to optimize their data processing and analysis pipeline. Each algorithm is explained with its fundamental principles, their application in neuroscience for cell assembly detection, and illustrated with published studies. Recognizing the similarities, advantages, and drawbacks of diverse methodologies may pave the way for developing new procedures for cell assembly identification to facilitate future endeavors in the understanding of brain activity.**

## INTRODUCTION

In 1949, Hebb [1] introduced the term “cell assembly” to define a group of neurons that are frequently coactive, thereby establishing strong interconnections, and acting together as a functional unit which could underlie cognitive function [2]. Cell assemblies are often regarded as representational entities, combining the behavioral correlates of individual neurons, perhaps binding together diverse features into an element for cognitive processing, e.g. a percept [3]. Some experimental questions also examine the occurrence of activations of assembly neurons in specific sequences, which could represent events in a temporal order. Complementarily, assemblies can be studied in terms of how their activations are processed by downstream targets [4, 5], where a cell assembly is defined by the activity it elicits. As in the case of binding, cell assemblies can span multiple brain areas [6]. Note also that assemblies may be found at different time scales depending upon the brain system studied [7].

Currently, two principal experimental approaches provide multiple neuron activation data at different time scales. In-vivo and in-vitro recordings with multiple microelectrodes typically resolve single unit activity on the order of milliseconds. Alternatively, calcium imaging allows recording large groups of neurons, and hence cell assemblies with known spatial locations, over long periods of time [8, 9]. Moreover, optical tools also provides the ability to stimulate cell assemblies [10]. However, the duration of calcium transients is on the order of hundreds of milliseconds [11, 12], resulting in slower temporal resolution compared to microelectrode recordings. Additionally, with current methods, only 10-20% of individual action potentials can be detected [13], with calcium transients mostly signaling when a given neuron is bursting. This limits the possibility to detect cell assembly activations when individual neurons fire only one action potential. As such, identified cell assemblies from calcium imaging data mostly consist of groups of neurons that burst together on timescales of around one second [14, 15].

Various methods have been devised to detect cell assemblies from microelectrode recordings and calcium imaging data, based on a range of theoretical and methodological frameworks. However, several existing techniques have only been tested on simulated recordings or have not been utilized outside of the laboratory where they were developed, and we hope that this review will encourage future studies to remedy this. Here, our focus is on methods that have been used to detect cell assemblies in in-vivo extracellular electrophysiological recordings from multiple microelectrodes, and that have been employed by researchers other than their developers. For recent reviews of methods used for detecting cell assemblies in calcium imaging, see [14, 16].

Early assembly detection methods were focused on event-based activity, identifying groups of neurons with activity correlated to specific behavioral variables, such as reward delivery, the animal’s position in space (e.g. [17, 18]), and sensory stimuli, potentially inducing bias. More recently, there has been increased interest in methods capable of detecting cell assemblies independently of environmental events, which can be particularly useful for examining brain activity without clear behavioral correlates. None of the approaches described here require information about the behavioral correlates of the neurons.

This review will provide an overview of several methodologies. Some techniques have the explicit ability to identify specific groups of recorded neurons as participating in cell assemblies [7, 19, 20], while other methods identify components or patterns for which all recorded neurons participate to some extent [21–24]. Furthermore, while some techniques are limited to simply detecting the co-activation of multiple cells within a given time window, others can detect cell assemblies as patterns of sequential activations of cells [7, 23, 24]. The subject of detection of sequential activations is beyond the scope of this review, and is only presented when the method detects assemblies as well.

Most techniques begin with the detection of co-activation of single pairs of neurons. This can then be expressed as co-activation matrices (such as correlation, similarity, or functional connectivity matrices), which are then used to extract more complex, multi-cellular, functional relationships. Three principal approaches will be discussed in separate sections below: dimensionality reduction (principal component analysis, independent component analysis, non-negative matrix factorization), network analysis (community detection, Markov processes), and agglomerative construction. We discuss the applications of each method as well as their strengths and weaknesses. This review aims to help investigators compare and choose among different protocols to find the best pipeline to detect and characterize cell assemblies in their data. Example publications are cited to provide more details useful for applying these methods as well as examples of some of the types of results that they can yield.

## DETECTING CELL ASSEMBLIES WITH DIMENSIONALITY REDUCTION

Dimensionality reduction maps high dimensional data into a lower dimensional space, and is commonly employed for analyzing large-scale data [25]. In neuroscience, an example of high-dimensional space is one formed by the activity of a neural population. Each neuron represents one dimension within this space. As such, a recording of 100 neurons over time would represent a 100-dimensional space and can be reduced to a smaller space of, for instance, 10 dimensions (or factors). The resulting 10 factors aim to preserve most of the information about neuronal co-activity in the original recording. In essence, each dimension in the new reduced space represents a recurrent pattern of population activity in the original (high-dimensional) spike matrix.

For linear dimensionality reduction techniques, each new dimension in the reduced space corresponds to a linear combination of all neurons, with each neuron having a weight quantifying its contribution to that factor. The interpretation of the obtained population pattern depends on the dimensionality reduction technique employed. For instance, if a technique is applied to a correlation matrix computed with Pearson’s pairwise correlations [26], multiple cells sharing high weights within a given factor would indicate that they have a high Pearson’s correlation score. The following section outlines the most popular dimensionality reduction techniques applied in the context of cell assemblies’ detection: principal component analysis, independent component analysis, and non-negative matrix factorization.

### Principal Component Analysis (PCA)

#### Basic principle

PCA is likely the most widely used linear technique for dimensionality reduction in neuroscience. Beyond neurophysiology, it is also useful to reduce high dimensional behavioral datasets (e.g. [27]). It reduces the original dataset into a set of dimensions (or factors) defined by orthogonal principal components (or eigenvectors, the cell assemblies), each resulting from the linear combination of the original variables (i.e., neurons) [28, 29]. Each component is associated with an eigenvalue that relates to the proportion of variance in the original dataset explained by that component. PCA is typically applied to a smoothed spike matrix representing binned spike counts (equivalent to firing rates), which are z-transformed. The first principal component explains the largest amount of variance (i.e., in co-activity of neurons), with subsequent components being necessarily orthogonal to the preceding ones, and explaining as much of the remaining variance as possible. By definition, there are as many principal components as variables (neurons) in the original dataset, but only a subset of them is needed to explain most of the variance.

#### Detection of cell assemblies with PCA

PCA is effective in detecting groups of neurons with correlated activity. Initially used in neuroscience to compress and synthesize neuronal responses across multiple trials [30–33], it was later employed by Nicolelis and colleagues [34, 35] as an indirect representation of cell assembly activity. In its most common implementation, as developed by Peyrache et al. [21], the algorithm starts by constructing a pairwise correlation matrix (**Figure 1A,B**) from the original spike matrix using Pearson’s pairwise correlation, a linear correlation measure that is normalized to the firing rates of the cells. The pairwise correlation matrix is then factorized into a set of orthogonal eigenvectors, each associated with an eigenvalue. Peyrache et al. [21] identified those principal components that explain a supra-threshold proportion of variance (termed “significant components”) using a threshold computed with the Marčenko-Pastur distribution [36]. Indeed, for a random normalized matrix, it has been demonstrated that eigenvalues follow a well-defined probability function, and are limited in magnitude by an upper bound [21]. The eigenvectors whose eigenvalues exceed this upper bound (**Figure 1C**) are considered to represent cell assembly patterns (**Figure 1D**). It remains to be proven whether this approach for random normalized matrices is completely applicable to neural data. Alternative approaches to detect significant components have been proposed [19, 22].

**Figure 1.**
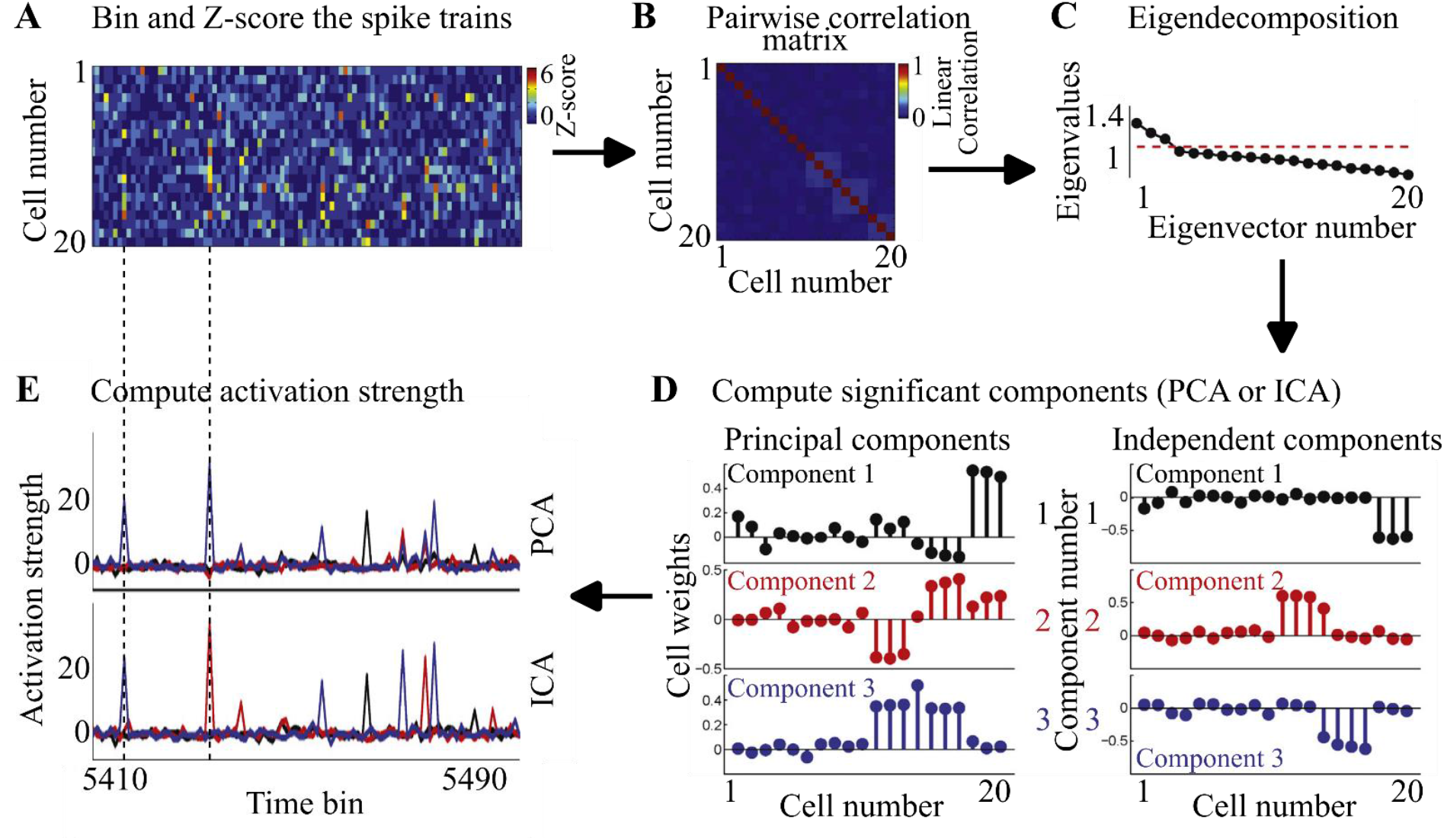
PCA and ICA implementation. Example with synthetic data. (Figure adapted from [22]). **A**. The data set (a fragment of which is shown here) composed of N (here 20) cells is binned in a window *dt* set by the user, and then firing is Z-scored. **B.** A pairwise correlation matrix is created, describing the co-activity of all pairs of cells in the dataset. **C.** The eigenvectors and eigenvalues are then extracted. The M (here, three) eigenvectors that have eigenvalues greater than a threshold (determined using the Marčenko-Pastur distribution, red dashed line) are considered as assemblies. **D.** Principal components and independent components obtained from the correlation matrix in B (see Table 1 for source code link). **E.** Activation strength of principal and independent components in the recording of A. Color codes are as in D.

**Table 1.**
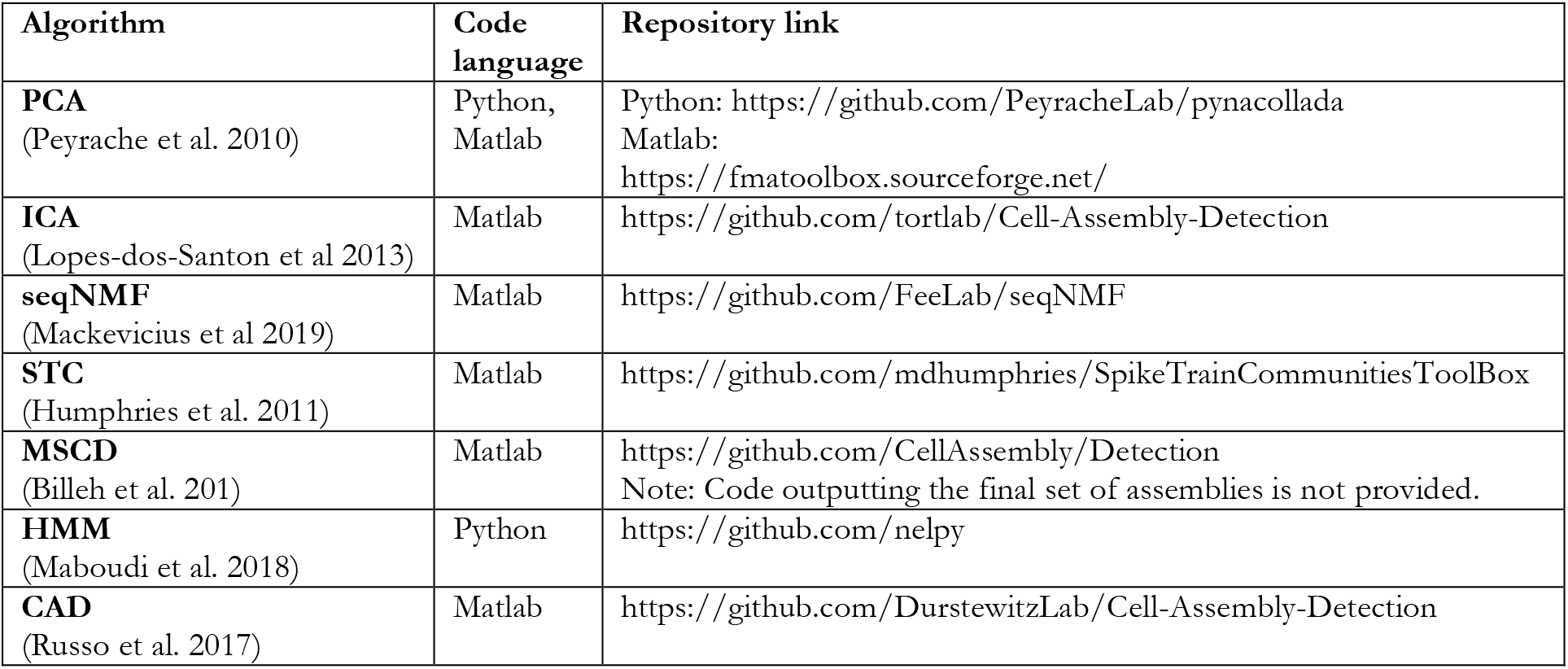
Availability of code for the cell assembly detection methods discussed here.

Peyrache et al. [37] used this method to detect cell assemblies in simultaneous recordings of dozens of prefrontal cortex neurons in behaving rats learning rules in a Y-maze. In this, and a subsequent paper [38], the authors identified cell assemblies in the medial prefrontal cortex that emerged upon learning, and were activated at the decision point of the maze. These cell assemblies were then replayed during slow wave sleep, synchronously with hippocampal sharp wave ripples. This was interpreted as supporting a role for coordinated hippocampal-prefrontal activity in memory consolidation. Since then, others have used this approach for cell assembly detection [e.g. 23–28]. Sjulson et al. [45] used PCA to detect cell assemblies and the directionality of information flow between the hippocampus and the medial striatum. They found that medial striatal cell assemblies decode spatial information from the hippocampus, and that conditioning to a cocaine stimulus increases the coupling of assemblies between these two structures.

#### Advantages and drawbacks of PCA

The main advantages of PCA are its operating speed, ease of use, and scalability up to very large datasets, even for thousands of neurons and tens of hours of recording. Computing the correlation matrix as well as matrix factorization are rapid operations, and do not require the user to provide values for multiple parameters. The only parameter that needs to be set by the user is the width of the binning window for detecting population patterns.

However, PCA shares some drawbacks with other dimensionality reduction algorithms, and these will be discussed in section 2.4. A drawback specific to PCA is the orthogonality between principal components. While the first component is optimized based on the direction of the largest variance in the multidimensional spike activity space, subsequent components are constrained to be orthogonal to this. Thus, a single neuron generally does not have a high weight in more than one assembly. This might not reflect the most optimal segregation of groups of correlated cells [46], leading to a lower probability of attributing appropriate weights to neurons participating to multiple cell assemblies [22].

### Independent component analysis ICA

#### Basic principle

Independent component analysis (ICA) has gained prominence as a data analysis technique in neuroscience [47]. Initially developed to segregate mixed sources into linearly additive subcomponents [48], ICA addresses challenges like the "cocktail party problem", where individual conversations must be distinguished from a recording in a noisy room. Similarly, in neuronal recordings, neuronal activity is comprised of a set of assemblies, and the objective is to reconstruct the identity of individual cell assemblies from the amalgam of activity from multiple cell assemblies and background noise. The Central Limit Theorem is the underlying basis of ICA. It stipulates that the result of combining two independent random variables is more Gaussian than the original variables. ICA improves the independence of components by rotating them and optimizing the non-Gaussianity of their combinations. In contrast with PCA, this results in non-orthogonal components.

#### Detection of cell assemblies with ICA

ICA was first introduced in neuroscience to identify groups of neurons exhibiting correlated firing activity [49]. The algorithm proposed by Lopes-dos-Santos and colleagues has become a widely used method for detecting cell assemblies [e.g. 3, 4, 41–47, 33–40]. This typical implementation of ICA for cell assembly detection involves a two-step procedure [22]. The algorithm first begins by reducing the dataset via PCA to yield significant principal components as above, and then uses ICA to rotate these components, yielding independent components.

Using this approach, Trouche et al. [52] identified spatially tuned cell assemblies in electrophysiological recordings. These assemblies encoded spatial information, and optogenetic inhibition of these cell assemblies impaired behavior during a spatial memory test. Guan et al. [65] utilized ICA to detect cell assemblies in bilateral recordings in the CA1 region of the hippocampus, identifying some assemblies restricted to either the left or the right hippocampus, while others extended bilaterally.

#### Advantages and drawbacks of ICA

Since the components in ICA are independent and not orthogonal, ICA is more suitable than PCA for including the same high-weight neuron in more than one cell assembly. Moreover, ICA inherits PCA’s advantageous traits: it is fast, scalable to large datasets, and user-friendly with only one parameter to set (the width of the binning window).

However, ICA comes with its own set of drawbacks. First, it assumes that the distribution of underlying variables is non-Gaussian, as components are retained when their non-Gaussianity is maximized. This assumption may limit the applicability of ICA in cases where the data distribution is Gaussian. Furthermore, due to the iterative nature of the optimization process, ICA is sensitive to initialization parameters and is susceptible to converging to local minima [66], i.e., the best result within a certain region of the solution space, but not necessarily the overall best solution. Therefore, repeated applications of ICA can return different results, and thus, different sets of cell assemblies. The effects of convergence to local minima can be minimized by always initializing the vectors to the same value to yield the same set of cell assemblies between runs. However, this will not ensure that ICA converges to its best solution. More intricate algorithms may offer a higher likelihood of finding the global optimum solution [67]. Additionally, different ICA algorithms can yield significantly different results based on their implementations [68], introducing variability that researchers must consider when interpreting findings.

A limitation shared by ICA and PCA is the arbitrary sign of highly weighted cells within a component. Highly positive or negative variables within a component can both indicate that cells are highly correlated [21]. It is crucial to note that the presence of both highly positive and negative weighted cells within a single component doesn’t necessarily represent two groups of anti-correlated neurons, but rather, this may result from poor separation of two assemblies [5, 6].

### Non-negative matrix factorization (NMF)

#### Basic principle

Non-negative matrix factorization (NMF) is a method designed to extract sparse features from high-dimensional datasets [69, 70], wherein each component represents groups of variables that are correlated. NMF decomposes the initial dataset into two lower-rank matrices, such that multiplying them together yields a result approximately equal to the initial dataset (**Figure 2A**). To achieve this approximation, values in the lower-rank matrices are iteratively updated to minimize the approximation error between their product and the initial dataset. The process terminates when the error converges −changes in the matrices become negligible− or when a user-preset maximum iteration limit is reached. Notably, NMF matrices consist exclusively of positive elements, facilitating interpretation of the resulting lower-dimensional matrices.

**Figure 2.**
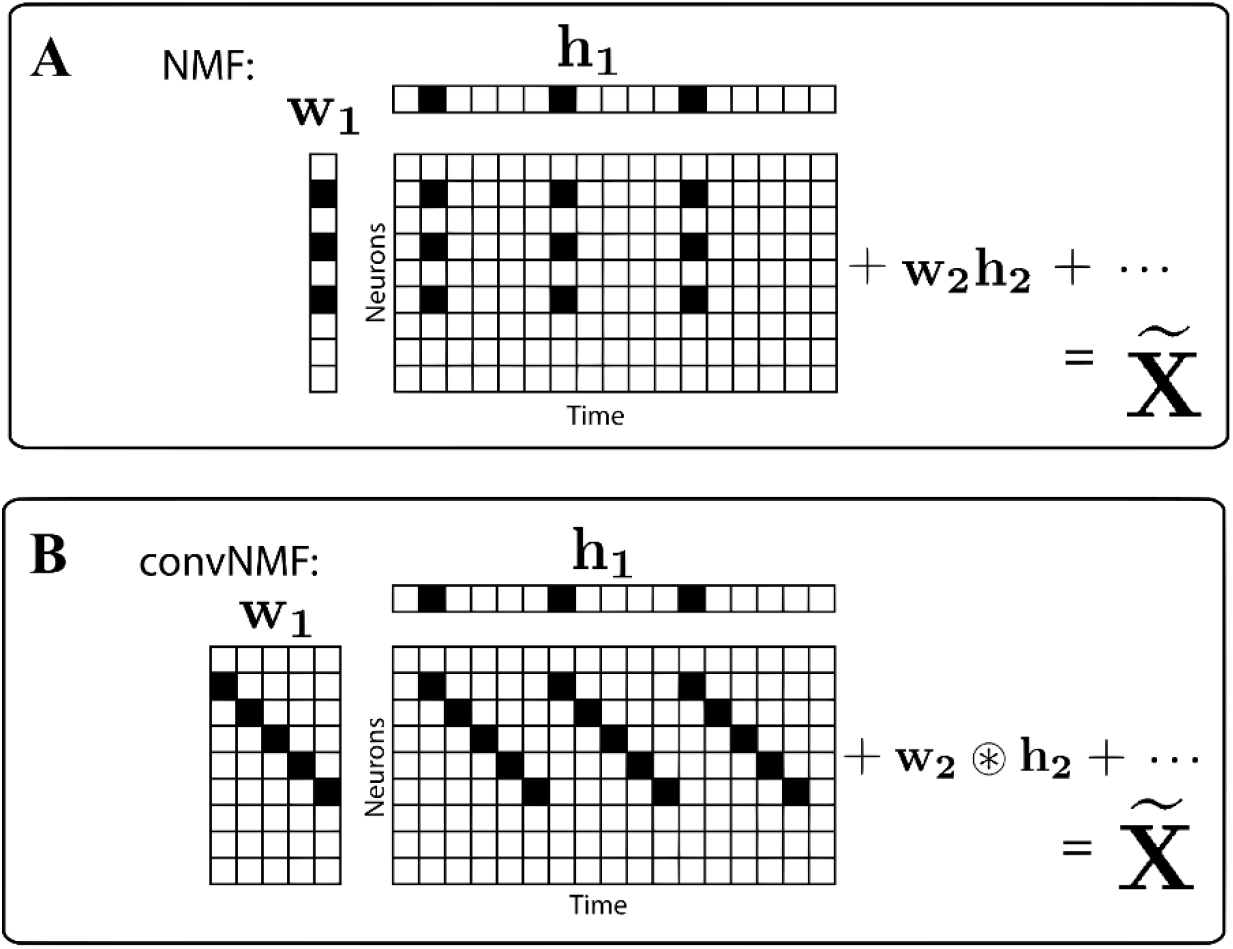
NMF and convNMF algorithms (adapted from [24]). **A.** NMF seeks to approximate a data matrix X composed of N neurons using two matrices, W and H. W is a matrix with N rows and K columns. N corresponds to the number of recorded neurons while K is set by the user and corresponds to the number of cell assemblies. H is a matrix with K rows and T columns, where T is the number of time bins in the data matrix. Matrices W and H can be decomposed into vectors w_1_…w_k_ and h_1_…h_k_. The goal is for W x H to be equal to the initial matrix. Every vector in W represents an assembly pattern (which cells are highly active in the assembly), and every vector in H represents a temporal pattern (the bins where the cell assembly is highly active). **B.** In convNMF, W is composed of matrices w_1_…w_k_, and each matrix stores a sequential pattern of neuron activations.

#### Detection of cell assemblies with NMF

In its application for cell assembly detection, NMF is employed on a binned spike matrix containing only positive values. It produces two lower-rank matrices of size K, where K denotes the number of patterns in the underlying dataset, typically chosen by the user. The first matrix denotes the identity of cell assemblies, with each vector representing a cell assembly (**Figure 2A**). The second matrix denotes assembly activations over time, where each vector represents moments when the corresponding cell assembly is active (**Figure 2A**).

With convolutional kernels, the algorithm can detect sequences of firing, capturing cell assembly activations with temporal structure across multiple time bins (**Figure 2B**, [24, 71]). Tingley and Buzsáki [72] applied this method to identify patterns of sequential cell activations in the hippocampus during sharp-wave ripple events. These assemblies were then used to assess whether neuronal activity in the lateral septum was related to the temporal organization of neuronal activations within sharp-wave ripples, and revealed a correlation with the magnitude of conjunctive activity rather than the specific order of cellular activity. See Terada et al. [69] for another application of NMF to detect sequencing in cell assemblies.

#### Advantages and drawbacks of NMF

The convolutional NMF algorithm offers the advantage of detecting assemblies with only positive weights, simplifying interpretation of results compared to PCA and ICA. Additionally, it demonstrates the capability to identify specific sequences of neuron activations, providing a nuanced understanding of temporal patterns within cell assemblies.

However, this method is not without its limitations. Similar to ICA, the results of NMF may vary between runs, and are contingent on the initialization of the lower-ranked matrices. Convolutional NMF can exhibit sensitivity to noise, potentially yielding correlated factors, or representing individual neuron activity through overfitting of the dataset [73]. To mitigate this issue, a regularization algorithm is often implemented [74, 75]. In terms of computational efficiency, convolutional NMF is slower compared to PCA and ICA, requiring more time to extract cell assembly patterns. Moreover, in the implementation by Mackevicius et al. [24], the method involves numerous parameters that necessitate manual tuning. These parameters include the number of cell assemblies, bin size, sequence length, as well as regularization parameters, and the learning rate of the optimization algorithm [24, 71]. The necessity for the user to pre-set the number of assemblies to find in the recording is a major limitation, since in most cases it is impossible to know how many assemblies are to be found. Manually adapting these parameters to a dataset can be intricate. It is worth noting that the authors provide tools in their toolboxes for automatic estimation of these parameters through recursive runs of the algorithm on the dataset, choosing those that yield the best results. However, this automated approach contributes to slower implementation.

### Summary: Advantages and disadvantages of dimensionality reduction for cell assembly detection

Dimensionality reduction algorithms exhibit notable advantages, first of all their relatively rapid run times. These approaches are particularly effective in detecting recurrent and stereotypical neuronal activation patterns, as evidenced in instances like bird songs [24, 76], however, they also share some limitations. One common challenge is the necessity to create a binned spike matrix, introducing edge effects where millisecond-scale changes in firing times may misattribute an action potential to a preceding or succeeding bin. To address this, datasets are often binned in overlapping intervals or smoothed using algorithms to distribute neuronal activity across multiple bins, mitigating edge effects [77–79]. Nevertheless, this can extend algorithm run time and potentially introduce duplicate patterns between bins, biasing cell assembly detection.

Dimensionality reduction methods factorize the dataset into a smaller number of variables. In the case of PCA and ICA, the number of components that can be obtained is limited to the number of cells in that dataset. This number is then further reduced when only the components explaining significant amounts of variance are retained. However, according to the concept of cell assemblies as fundamental computational blocks of brain function [1] the number of cell assemblies (i.e., combinations of co-active neurons) is not necessarily so limited. On the contrary, one can argue that nervous system processing might benefit from more cell assemblies than individual neurons. Indeed, if the fundamental computational blocks were individual cells, then the brain capability would be limited by their overall number. Instead, if the basic computational units were all the possible combinations of neurons, this would extend computational capabilities multifold. Therefore, in a given recording of a limited sample of neurons, it is possible that the number of cell assemblies exceeds the number of cells. Dimensionality reduction methods would be unable to detect all these patterns as independent cell assemblies. Additionally, results are presented as continuous weights, representing each cell’s contribution to a component. Consequently, all neurons contribute to each detected assembly, and the exact identity of cells participating in a cell assembly is not provided. To overcome this limitation, threshold measures have been proposed to identify those cells most involved in specific cell assemblies [5, 6, 59, 60, 63, 64, 80]. Finally, note that, like other approaches, no distinction is made between co-activations related to functional relations between the neurons, or from common driving from another source.

## CELL ASSEMBLY DETECTION WITH NETWORK ANALYSIS

Beyond dimensionality reduction, other methodological approaches have been proposed to detect cell assemblies in high density neural recordings using a variety of statistical or analytical approaches. Some of them are based on techniques developed to analyze complex networks.

### Community detection

#### Basic principle

Community detection is a network analysis method designed to examine complex graph structures. A graph consists of nodes interconnected by edges, where edges may carry numerical weights representing relationships between nodes. Networks often exhibit communities: subsets of nodes with stronger internal connections and weaker connections to other parts of the graph. Originally devised for social networks and biological ecosystems [81], community detection algorithms have been applied in various fields with graphs containing highly connected networks separated by loosely connected nodes.

Community detection has also been proposed for the detection of cell assemblies in neurophysiological recordings [19, 20]. In this case, the nodes correspond to neurons within the dataset. Between two nodes, the edge weights correspond to the degree of co-activation of the two neurons. Detected communities would represent groups of highly connected neurons in the graph. It is important to note that with standard community detection methods, each node can only belong to a single community. Thus, in implementation to neuronal data, each neuron can only belong to a single cell assembly.

#### Detecting cell assemblies with Spike Train Communities (STC)

Humphries [19] introduced Spike Train Communities (STC), a community detection-based method for finding cell assemblies. The algorithm begins by creating a pairwise similarity matrix based on cosine similarity of spike trains at a user-defined timescale (**Figure 3A-B**). The cosine similarity is influenced by the pattern of neuronal activations and the relative activity of each cell within the population vector, but not the scale (or absolute value) of that activity. The obtained pairwise similarity matrix is then factorized with eigen-decomposition, yielding a set of eigenvectors and their corresponding eigenvalues. Only the M eigenvectors that have positive eigenvalues are retained (**Figure 3C**). These eigenvectors are used to define a set of coordinates within an M-dimensional eigenspace. This is then used to cluster the data into groups of highly correlated neurons via k-means (**Figure 3D**). On each iteration, k-means detects a different number of clusters, numbered between 2 and M+1 (**Figure 3E**). The optimal number of clusters can be computed as those communities that maximize within-cluster connections and minimize between-cluster connections (**Figure 3E**). In addition to detecting cell assemblies, this method has been used to identify similar spike trains fired by a single neuron across multiple trials [61, 82–84].

**Figure 3.**
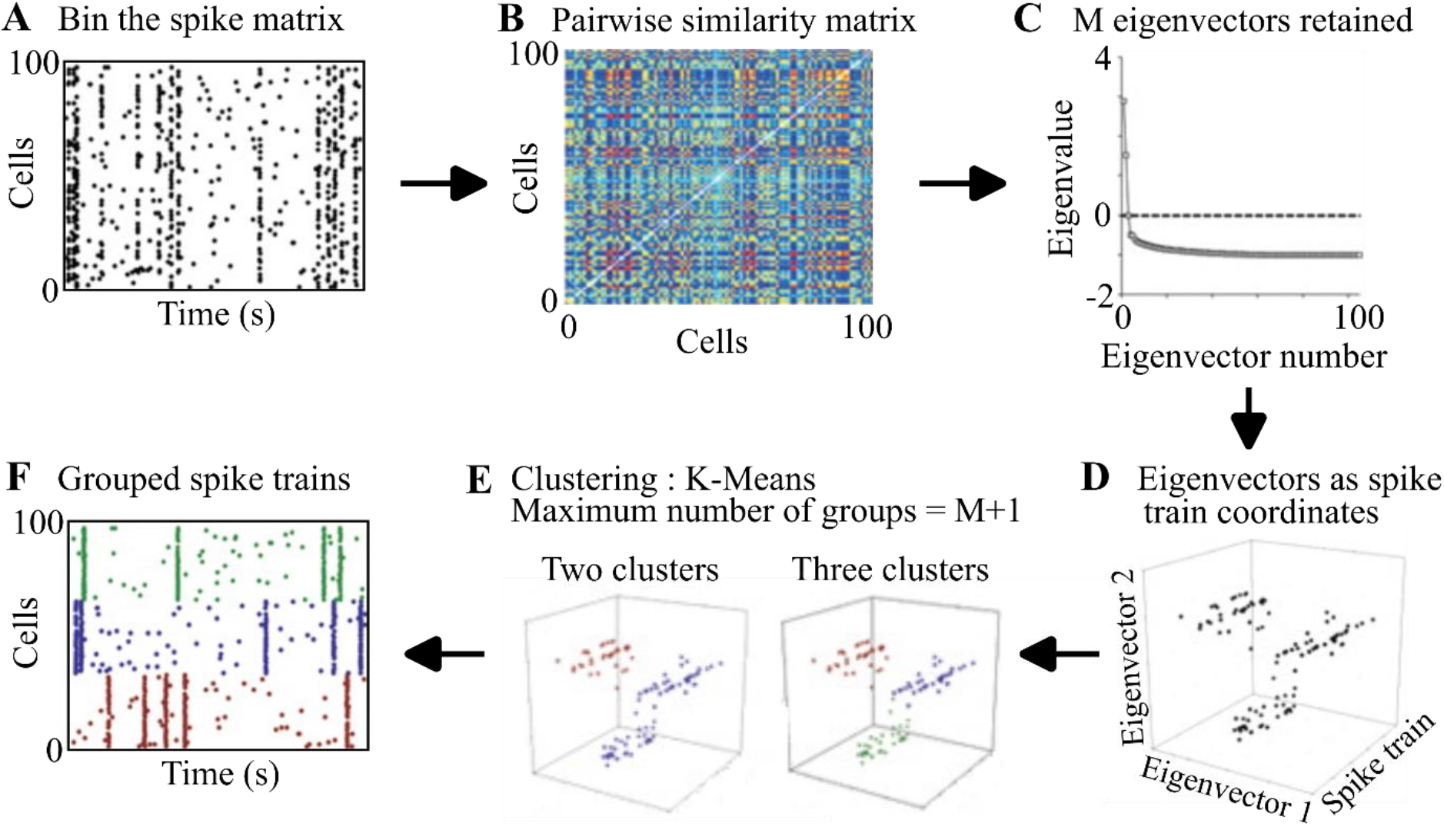
Implementing STC (figure adapted from [19]). **A.** The dataset composed of N (here 100) cells is binned into windows *dt*, a value set by the user. **B.** A pairwise similarity matrix is created describing the co-activity of all pairs of cells in the dataset. Red indicates highly correlated cell pairs, while deep blue are anti-correlated pairs **C.** The eigenvectors and eigenvalues are then computed. The M eigenvectors that have positive eigenvalues are retained. **D.** Spike trains are represented in the coordinates composed of the M eigenvectors with the highest eigenvalues. The eigenvectors are each N-entries long. Then, each eigenvector defines a set of coordinates in an M-dimensional space for every spike train. Within this eigenspace, the closer two spike trains are, the more similar are their activities. **E.** Clustering is carried out on the spike trains using k-means, iteratively detecting communities k = 2…, M. For each k, a metric based on the number of links within the communities is computed to determine how well the clustering fits the underlying dataset. The k with the best fit to the data is retained. **F.** The spike trains from A, grouped according to the communities they belong to. Colored vertical lines correspond to cell assembly activations.

McMahon et al. [84] applied this method to recordings of spinal cord interneurons in cats performing an air stepping test. They identified two cell assemblies in which interneurons participate. This allowed the authors to assess the anatomical location of cells participating in these cell assemblies, finding no correlation between a given cell’s location along the rostro-caudal axis of the spinal cord and its participation in one or the other cell assemblies. They also found that activations of these two cell assemblies were respectively correlated to flexor or extensor muscle bursts during stepping.

##### Advantages and disadvantages of STC

Similarly to PCA and ICA, this method does not require the user to set any parameters other than the timescale at which assemblies will be detected. STC can be implemented in a binless version, centered on spikes rather than time bins, yielding superior results, but slowing computation by an order of magnitude. Note also that k-means is sensitive to its initialization parameters [85], and could converge to local minima across different runs, resulting in different clusters each time, and yielding a set of non-optimal clusters. Humphries takes these limitations of k-means into account, and employs methods to minimize these problems. Another disadvantage is that a single neuron cannot participate in more than one assembly.

### Using Markov chains to detect communities

#### Basic principle

Markov processes are stochastic models that infer the probabilities of sequences of states in a system. Each state corresponds to an observable variable or event. The future state of the system depends only on the current state, the current connections within the graph, or in the case of a neuronal recording, the pairwise correlation metric.

In the context of network analysis, Markov processes offer a powerful tool for identifying communities within a given graph. The fundamental idea involves assessing interactions between nodes to pinpoint groups that are densely connected and closely associated [86]. This is done by examining how a process that evolves with time interacts with the Markov process. Schematically, this can be represented by a person randomly walking within the graph. More highly connected parts of the graph would trap the random walk process for longer periods of time compared to parts that are loosely connected. Highly connected parts of the graph would correspond to communities. Within the graph, there can be communities of various sizes. If the process evolves within the graph for short periods of time (called “Markov Times”), the areas encompassed by the process would be small, preventing the detection of larger communities, which appear over longer times. However, long Markov Times might affect the detection of smaller communities.

In neuronal recordings, the events are spiking activity of neurons, and these highly connected compartments would represent cell assemblies. Communities detected at smaller Markov Times would detect many cell assemblies composed of small numbers of neurons, while conversely, longer Markov Times would detect fewer cell assemblies composed of larger numbers of neurons. Understanding the impact of Markov Time on community detection is crucial for analysis design and interpretation. The choice of Markov Time influences the granularity of identified cell assemblies, offering flexibility to adapt the analysis based on the desired level of detail or comprehensiveness.

##### Markov Stability Community Detection (MSCD) to detect cell assemblies

To detect cell assemblies, Billeh et al. [20] proposed a method employing Markov Stability for Community Detection (MSCD). First, the algorithm constructs a pairwise coactivity matrix by computing the so-called “functional connectivity” between neurons (note that this does not necessarily correspond to anatomical connectivity). The authors define this metric as the expected contribution of one neuron’s activity to that of another neuron. The algorithm proceeds by constructing a template for each action potential fired by a neuron over time. The functional connectivity between two neurons A and B is determined as a function of the likelihood that neuron B fires shortly after neuron A. Excitatory connections are modeled as activation that decays exponentially with time (**Figure 4A**). Inhibitory connections are modeled using an inverted exponential profile (**Figure 4B**). This process is repeated for all pairs of neurons in the dataset, resulting in a pairwise functional connectivity matrix. Given the asymmetry of relationships between neurons, the matrix values can be likened to directional connections in a graph.

**Figure 4.**
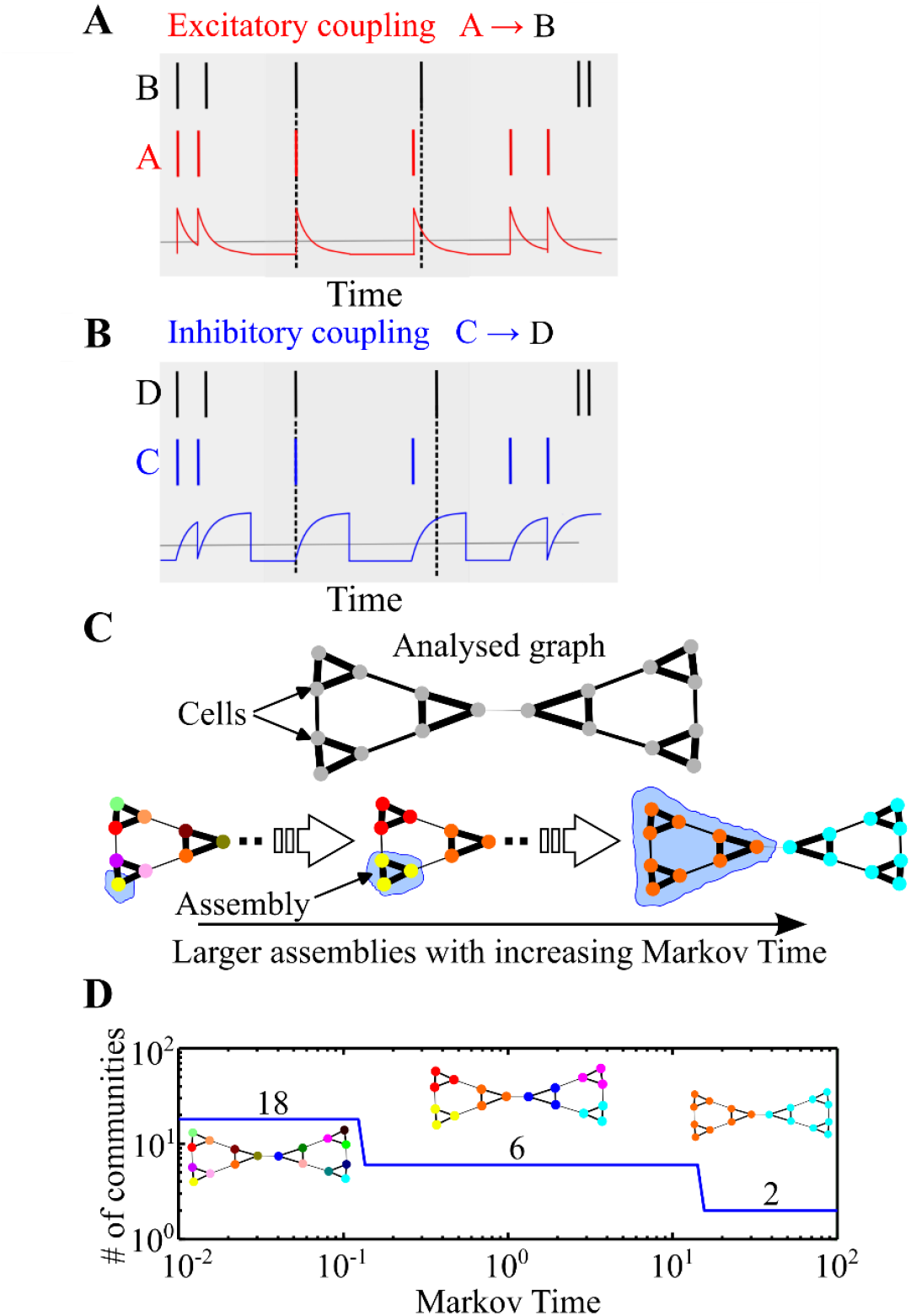
Schematic representation of the MSCD method (adapted from [20]). **A.** Computation of functional connectivity between pairs of neurons. Predictability of neuron A for neuron B is modeled as a positive prediction that decays exponentially with time. The top two lines are spike trains of the neurons. The third trace is the association function that governs the degree to which spikes of B are taken to be predicted by A. This coupling is highest when neuron B fires immediately after neuron A (left vertical dashed line) but is reduced for longer delays (about 50% for the right dotted vertical line). **B.** Negative coupling between neuron C and neuron D is modeled using an inverse exponential profile. Note that when neuron D fires immediately after neuron C, the predictive association is nil (left vertical dotted line), but the negative predictive association is partial when neuron D fires at a greater delay after neuron C (right vertical dotted line). **C.** A process evolving within the associational connectivity graph can be used to detect communities. In this example graph, there are 18 nodes (neurons). Nodes are connected through edges. The width of the edge represents the strength of the functional connectivity between two nodes. At longer Markov Times, the process can explore larger parts of the graph, yielding larger cell assemblies. At each Markov Time, the nodes of a given color represent a different community. **D.** The number of detected communities is plotted as a function of Markov Time. At very brief Markov Times, 18 communities are detected, corresponding to the number of cells in the dataset. At longer Markov Times, 6 communities are detected, corresponding the small triangular structures, each of which is composed of three highly connected cells. At even longer Markov Times, 2 communities are detected, each corresponding to nine cells that are strongly connected.

Communities within this graph are then identified using a technique based on Markov processes (**Figure 4C**). This can yield a large number of communities detected across multiple Markov Times (**Figure 4D**). Interestingly, some communities are only detectable at specific Markov Times and may not fully represent the community structure within the graph. To identify consistently detected cell assemblies across Markov Times, the authors propose two measures of robustness, considering the most consistently detected communities as the most meaningful. This algorithm was implemented by Miyawaki et al. [87]. Gonzalez et al. [15] also employed a method based on this algorithm to detect cell assemblies in calcium imaging recordings.

##### Advantages and disadvantages of MSCD

One strength of this algorithm, relevant for certain experimental questions, stems from the directionality of the computed functional connectivity matrix. Unlike the similarity matrices described earlier, the relationship between neurons here is asymmetrical and dependent on precise spike times. For example, the predictability of neuron A’s spike for those of neuron B might not be equal to predictions in the opposite direction. Furthermore, the user can specify which neurons are excitatory and which ones are inhibitory, and the influence of their activations is modeled differently (Figures 4A and 4B, respectively). Also, the exact identity of cells composing a cell assembly is known.

However, this algorithm is computationally more demanding than those described above. Moreover, the results can vary between runs since this community detection algorithm is sensitive to initialization parameters. Consequently, the community detection framework is typically executed between 100 and 500 times per Markov time to identify the most consistently detected cell assemblies and reduce dependence on initialization parameters. Additionally, at a given Markov Time, any given neuron can only belong to a single cell assembly, limiting the number of cell assemblies that can be detected in the dataset.

### Hidden Markov Models (HMM)

#### Basic principle

Hidden Markov Models (HMM) extend the concept of Markov processes, and early applications included speech recognition and DNA analysis. In the Markov processes discussed above, nodes represent the observable variables of the underlying dataset (neuronal activations). However, the interactions between neurons may be influenced by unmeasured variables in the original dataset, such as global dynamics like UP and DOWN states, ripple events, and cell assemblies. HMM introduces hidden nodes to represent these unmeasured variables, which are inferred from observable measures, and the two sets can interact. An observable variable can contribute to several hidden nodes with different weights, and the hidden nodes are generally expressed as linear combinations of the observable variables. The hidden Markov model computes the hidden variables that best explain the interactions between the observable variables.

#### Detection of cell assemblies with HMM

A few authors have proposed the use of HMM to detect cell assemblies [23, 88, 89]. Maboudi et al. [23] constructed a hidden Markov model using population burst events, defined as time windows where the number of spikes fired by the neuron population is at least three standard deviations greater than the mean over the entire recording. In all implementations, the user must select, as a parameter, the number of hidden nodes describing the unmeasured variables (here, cell assemblies) that characterize spiking activity relations between neurons in the dataset. A principal goal of the algorithm is to assign to each neuron a weight defining its contribution to each of the assemblies.

This method allowed Maboudi and colleagues [23] to model sequentially active cells forming assemblies from recordings in the hippocampus, and predict the positions of behaving rats from the place fields of the neurons. Furthermore, distinct sequences of replay in the hippocampus were identified without knowledge of the behavioral correlates of the individual neurons. The method described by Maboudi and colleagues was also implemented in other studies [50, 90, 91]. Noguchi et al. [90] used this method to detect sequential patterns in hippocampal neuronal activity during sharp-wave ripples. This allowed the authors to study how the activity of interneurons preceding sharp-wave ripples influenced the structure of subsequent neuronal sequences.

##### Advantages and disadvantages of HMM

This method allows a single neuron to participate in several cell assemblies (unlike community detection algorithms). Like the convolutional NMF algorithm, it can also detect patterns of sequential activations of cells. Furthermore, the number of cell assemblies that can be detected by this algorithm is theoretically unlimited and can exceed the number of neurons in the dataset.

A main limitation of this method is that the user needs to estimate how many cell assemblies are to be found in the recording, which is difficult. If the user overestimates the number of assemblies, this leads to the distribution of noise among the assemblies; the algorithm may seek to explain residual spike activity as cell assemblies. On the other hand, a low estimate will lead to missing some assemblies. Due to the higher number of user pre-set variables and the need to learn hidden variables from the observable dataset, a main drawback of hidden Markov models relative to other Markov process methods is the long computation time.

## AGGLOMERATIVE CONSTRUCTION OF GROUPS

### Basic principle

Instead of reducing dimensions or clustering the population of cells based on functional relationships between them, agglomerative algorithms start with small templates, such as a pair of co-active cells. The process then progressively enlarges the group by finding another cell that tends to be coactive with the existing group, creating a three-cell group. The algorithm continues this agglomerative construction iteratively, searching for neurons significantly co-active with an n-cell template to create an n+1 template. This agglomerative construction persists until no new neurons can be added to the template. Various statistical methods can be employed to assess neuronal co-activity and its significance.

### Agglomerative detection of cell assemblies with CAD

Russo and Durstewitz [7] developed the CAD (Cell Assembly Detection) algorithm in order to identify groups of cells that are active together above chance levels. The method first creates a binned spike matrix, and then tests all combinations of pairs of cells for significant co-activation (**Figure 5A-B**). For any pair of neurons, the number of spikes fired by the two neurons is counted at several time lags (time lag of 1 bin, 2 bins… etc), and the time lag with the largest number of coupled spikes is retained. The counts are then tested for significance (**Figure 5C**) and cell assemblies are progressively characterized (**Figure 5D-E**). A group of cells is considered as a cell assembly when no new cells can be added (**Figure 5F**).

**Figure 5.**
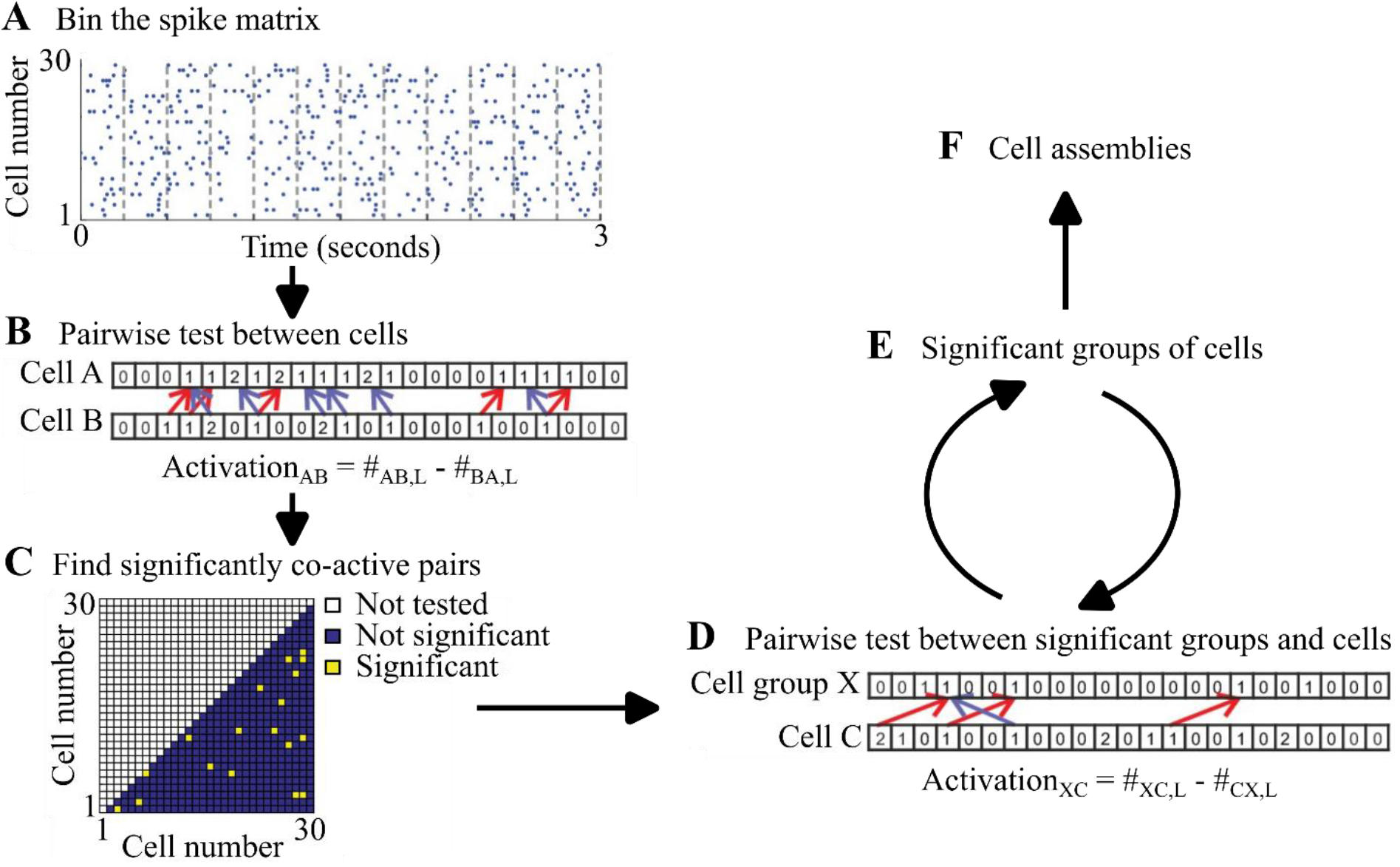
The agglomerative assembly detection of Russo and Durstewitz [adapted from 6]. **A.** The dataset composed of N cells is binned in a window *dt* set by the user. **B.** The co-activity of all pairs of cells in the dataset is assessed. #_AB,L_ is the number of times cell B is active after cell A at the time lag L, #_BA,L_ is the number of times cell A is active after cell B at the time lag L. Activation_AB_ is the difference between the two. **C.** The significantly co-active pairs of neurons are determined. **D-E.** Cell groups are progressively expanded. The pairwise test is performed between significant cell groups X found in the previous iteration and each remaining cell. Cells are added to the cell group if they are significantly co-active with the cell group. **F.** The cell assemblies are obtained when no new cells can be added to the cell groups.

In recordings, the number of spikes fired by a neuron can vary depending on the behavioral state of the animal, common driving inputs, and neural oscillations, giving rise to stationarities. To ensure that these global variations in cell firing rate do not influence the correlation statistics between cells, Russo and Durstewitz devised a novel pairwise test (**Figure 5B**). The algorithm measures the asymmetry between the spikes of two neurons (or a neuron and a group of cells), comparing the number of spikes in the spike train (“forward pass”) and the spike train replayed in reverse order. Indeed, if two cells A and B emit action potentials independently, it would be expected that the number of spikes fired by cell A after cell B to be equal to the number of spikes fired by cell B after cell A. If this number is skewed in one direction or another, it is inferred that the two cells have a functional relationship. This difference can then be used to compute the significance of the interaction between cells A and B.

This method was employed to detect cell assemblies in high-density electrophysiological recordings [91, 92]. Oettl et al. [92] analyzed midbrain dopaminergic and striatal projection neurons with this method, and found that striatal projection neuron activity preceded midbrain dopaminergic neuron activity. Furthermore, cell assembly activity changed during the trials, and was correlated to the value of a stimulus.

### Advantages and disadvantages of CAD

With agglomerative methods, the number of cell assemblies that can be detected is very high and could extend to the number of combinations among the recorded neurons. Furthermore, as cells are assessed individually for their participation in cell assemblies, the final identity of neurons composing each cell assembly is known. The same neuron can be included in more than one assembly. The method can also detect patterns of sequential activations of cells. Furthermore, the pairwise coactivity metric used by Russo and colleagues is robust against stationarities.

Note that, like many methods, the sensitivity of agglomerative clustering is dependent upon how often the cell assembly is active during the recording. Indeed, if a large cell assembly is active only a few times across a given recording there might not be enough statistical power to detect it as an assembly, but multiple fragments of it could be detected. One potential problem arises from the fact that when one neuron’s activity is evaluated to be considered as part of an entity, the activity of the latter is considered as a whole. Therefore, the neuron may be considered to be significantly co-active with the group only because it is significantly active with a subset of it.

## IMPLEMENTATION AND COMPARISON OF THE RESPECTIVE METHODS

For all algorithms described above, the availability of the code, as well as links to the official repositories are presented in **Table 1**. To help the user select the best method for their analysis, **Figure 6** summarizes their main features.

**Figure 6.**
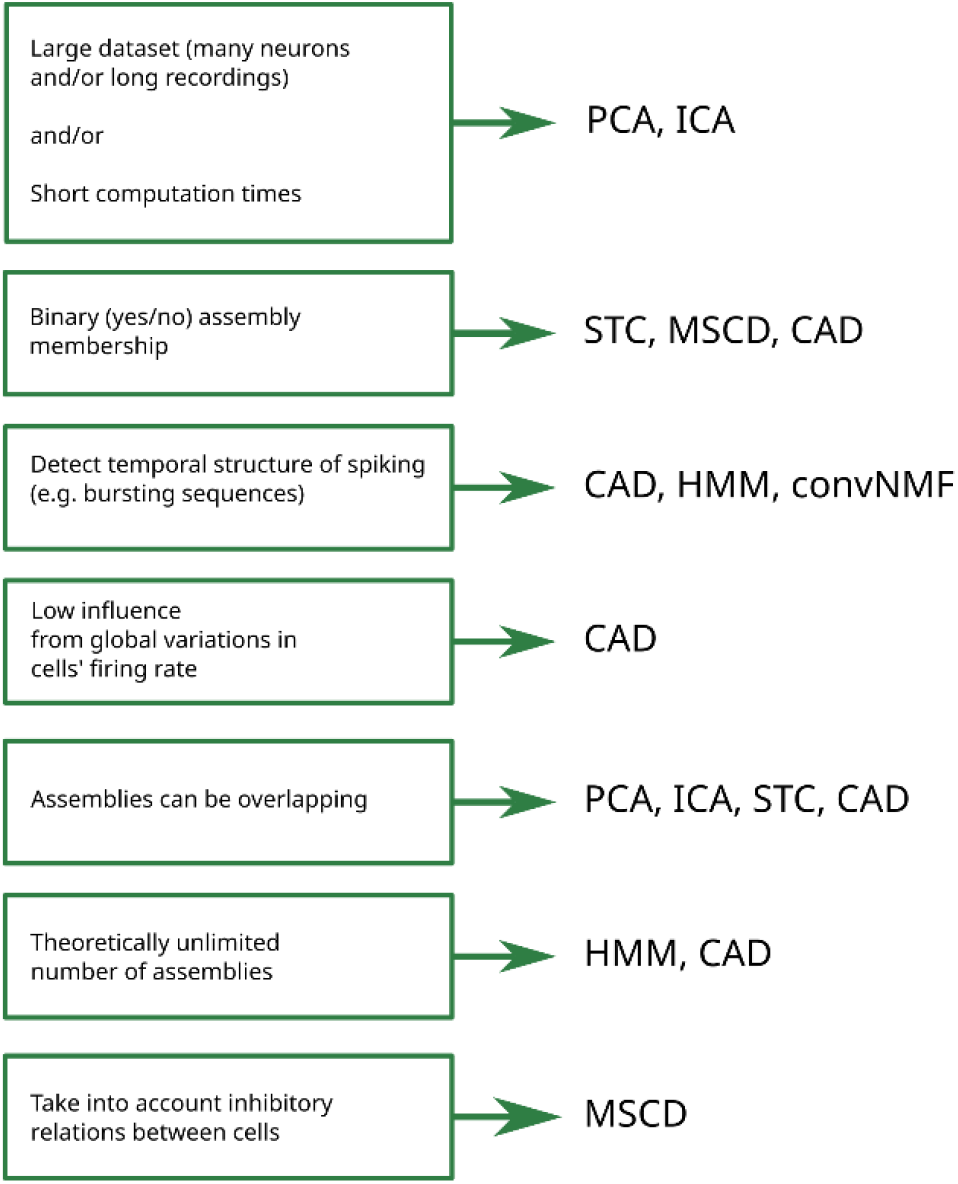
Decision chart for the selection of cell assembly algorithms based upon experimental aims.

For comparison, we implemented these methods on a simulated recording of 50 cells forming 50 cell assemblies over 60 minutes. Each assembly was composed of 3 cells (with individual cells participating in multiple assemblies). Cells were simulated using a Poisson point process with a mean firing rate of 1 spike per second. Cell assembly activations were added at random time points in the dataset and were active once per minute; all neurons composing the cell assembly fired together within a 30 ms window by adding random jitter to spike timings. Within cell assembly activations, the member neuronal activity was simulated with a Poisson point process with a mean firing rate of two spikes per assembly activation. The different methods were implemented using the toolboxes available online (**Table 1**). The code from Billeh et al. [20] was modified to run faster by reducing the binning window ten-fold, optimizing the thresholding function, and enabling parallel processing on multiple cores (these modifications only impacted algorithm run time without altering the result). Where necessary, a binned spike matrix was constructed using a 30 ms binning window and a 15 ms overlap between two successive windows. In the methods that required setting a time window for cell assembly detection, this was set to 30 ms. For the remaining parameters, the default settings were used for the various methods. The results are presented in **Table 2**. There are large variations in run times, with some methods completing the cell assembly detection task in only a few seconds (PCA, ICA), and other taking minutes or longer (MSCD, HMM, CAD). Note that none of the approaches detected the exact identity of the 50 cell assemblies present in the underlying dataset. Furthermore, the results also differed between methods and no two algorithms returned identical sets of cell assemblies.

**Table 2.** Performance and metrics of the methods used in this review. The data set was a simulated recording of 50 cells forming 50 cell assemblies over 60 minutes. Each assembly was composed of 3 cells. See text for the selected parameters for these implementations.

| Algorithm | Run time (seconds) | Number of assemblies detected | User-set parameters | Notes |
| --- | --- | --- | --- | --- |
| **PCA**<br>(Peyrache et al. 2010) | 0.4 | 18 | 1 | Parameter to set: Time bin duration |
| **ICA**<br>(Lopes-dos-Santon et al 2013) | 4.3 | 18 | 1 | Parameter to set: Time bin duration |
| **seqNMF**<br>(Mackevicius et al 2019) | 12.0 | (user set to 50 here) | 3-6 | Parameter to set: number of assemblies to detect (among others). Some parameters can be optimized with algorithms in the toolbox. |
| **STC**<br>(Humphries et al. 2011) | 12.0 | 10 | 1-2 | Parameters: Time bin and possibility to optionally choose the correlation method |
| **MSCD**<br>(Billeh et al. 2014) | 120.4 | 2 | 1 | Parameter: Markov Time. List of Markov Times must be provided in advance. |
| **HMM**<br>(Maboudi et al. 2018) | 97.4 | (user set to 50 here) | 2 | Parameters: Time bin; number of assemblies. |
| **CAD**<br>(Russo et al. 2017) | 2918.7 | 90 | 2-3 | Parameters: Time bin; time lag. Possibility to optionally choose the significance threshold. |

## DETECTION OF ASSEMBLY ACTIVATIONS

Some techniques can identify members of cell assemblies [7, 19, 20], while other methods identify components or patterns for which all recorded neurons participate to some extent [21–24] (but see [5, 6, 59] for methods to extract individual members from their components). After identifying cell assemblies, one may wish to detect the exact moments in time where the activations of a cell assembly occurred. This can be used to characterize behavioral correlates associated with these activations or the reactivation of cell assemblies during various behavioral and brain states such as sleep (e.g. [93]). Determining the occurrence and eventually the intensity of cell assemblies’ activations is a problem distinct from the identification of the cell assemblies.

Certain methods described here return the identity of cell assemblies, but not the times when these assemblies are active (PCA, ICA, STC, MSCD). In order to obtain the activations of an assembly, typically, matrix convolution is used. Matrix convolution involves the multiplication of a matrix with a kernel (a pattern) to extract relevant features from the former. The kernel is a pattern that can be represented by either a vector or a matrix, generally of smaller size than the data matrix. The convoluted resultant matrix is therefore weighted by the kernel.

To detect cell assemblies’ activations, matrix convolution is employed on spike matrices using vectors representing cell assemblies’ identities as kernels. These vectors contains either the binary identities of assembly members, assembly member or not, or continuous weights representing the degree of participation of the neuron to the given assembly. The vector resultant of the convolution represents the activation of the assembly over time. This will either be a discrete variable for assemblies with binary membership, or continuous for assemblies with weights for each recorded cell.

In the case of weighted vectors, to overcome the limits of the matrix convolution, Peyrache et al [21] introduced a new metric in order to compute the similarity of the population vector in the binned spike matrix (**Figure 1E**). The method computes the reactivation strength as the square of the weighted vector (with a nulled diagonal) projected onto the binned and z-scored spike matrix. Due to the squaring of the weighted vector and the exclusion of the diagonal, this method takes into account the co-activity of cells, rather than individual cells’ firing rates. The resulting reactivation strength can then be Z-scored and epochs which surpass a certain user-set Z threshold can be retained as significant reactivations of the assembly. By utilizing a weighted cell vector, this method takes into account population activity as a whole (silent as well as co-active neurons, and positively and negatively correlated neurons). Furthermore, reactivation strength is sensitive to the bursting behavior of cells. Epochs when the cell assembly displays bursting behavior will have higher reactivation strengths than others. However, using this method, it is difficult to control the number of cells whose co-activation is required in order to consider a specific epoch as being a reactivation of the cell assembly. Indeed, bursts of activity of a subset of cells in the assembly can be sufficient to reach significance. As such, it is possible that two non-overlapping sets of cells could result in detected activations of the “same” cell assembly.

## CONCLUSIONS AND PERSPECTIVES

Despite the diversity of the methods designed to detect cell assemblies using various mathematical approaches, they all share some common limitations.

It is important to note that in recordings of large nervous systems, such as in mammals, it is highly unlikely that any method can detect all members of a cell assembly, due to technical limitations in the number of cells that can be recorded simultaneously. Thus, an assembly consisting of only three members may still represent the “tip of the iceberg” which contains many other unrecorded members.

The cell assembly is not simply a static representation, but rather functions by transmitting a signal that may well be processed by other assemblies spanning the same, overlapping or different structures. Currently developed methods seek to detect cell assemblies by analyzing the activity of typically a single structure sending the signal (but see [6]).

The techniques presented here are best at detecting assemblies that recur fairly frequently in the dataset. However, some cell assemblies, perhaps involved in processing rare behavioral or cognitive events may occur only sporadically, and thus may not be reliably detected.

At this time, to identify cells within assemblies, no empirical evidence exists to justify the choice between either continuous weights or binary membership, and thus it is difficult to compare the accuracy of these respective methods. The choice of the method should pertain to the theoretical basis of the experimental question and the experimental design.

Synchrony between neurons can come from local connectivity, or one group of neurons exciting another, but also from mutual driving by a third structure, a sensory input, or another event such as UP and DOWN state transitions etc… There is a risk that neuronal correlation measures could be influenced by these events. Stationary spike trains arising due to such events can be reduced by detecting cell assemblies during a neutral period such as sleep (taking into account eventual sleep oscillations). Russo and Durstewitz [7] propose another method for dealing with stationarities.

Modern tools allow the recording of hundreds or even thousands of neurons simultaneously across different cerebral structures. The development of modern computational tools for the study of cerebral circuits enabled the exploration of population activity in novel ways including the detection of cell assemblies, which previously were only theoretically hypothesized. Some methods, such as those based on dimensionality reduction or clustering approaches, also simplify analysis of the dataset by reducing it into a fewer number of variables, while preserving its principal properties. Others can detect larger numbers of cell assemblies than the number of neurons in the original dataset, and thus may be able to more completely explore and assess the dynamics and interactions of cell assemblies. Furthermore, most algorithms are based on neuronal activity discretized into a binned spike matrix, and some evidence suggests that binless detection algorithms may return superior results [19]. Future algorithms could address this issue, as well as enable a more thorough view of the available cell assemblies through more sensitive algorithms as well as algorithms that are better optimized to run on very large datasets (e.g. [94–96]). Finally, in recent years, population activity has been investigated with the use of machine learning and neural networks (e.g. [97]) and such approaches could be employed for cell assembly detection.

## ACKNOWLEDGMENTS

We thank Ralitsa Todorova for constructive discussions and Noé Hamou and Vinicius Lima for useful comments on the manuscript. This work was funded by a LabEx MemoLife grant to SIW and MNP and by an FFRE grant to MNP. We apologize to the numerous authors inspiring this work, but whose studies we could not cite and acknowledge that the selection of references has left out many important contributions.

## AUTHOR CONTRIBUTIONS

Initial draft GM and MNP. Simulations and testing of methods: GM. Graphics and visualization: GM and MNP. Writing: GM, SIW, and MNP.

## References

1. Hebb DO (1949) The organization of behavior; a neuropsychological theory. Wiley, Oxford, England

2. Eichenbaum H (2018) Barlow versus Hebb: When is it time to abandon the notion of feature detectors and adopt the cell assembly as the unit of cognition? Neurosci Lett 680:88–93. 10.1016/j.neulet.2017.04.006

3. Gray CM, Konig P, Engel AK, Singer W (1989) Cortex Exhibit Inter-Columnar Global Stimulus Properties. Nature 338:334–337

4. Buzsáki G (2010) Neural Syntax: Cell Assemblies, Synapsembles, and Readers. Neuron 68:362–385. 10.1016/j.neuron.2010.09.023

5. Boucly CJ, Pompili MN, Todorova R, Leroux EM, Wiener SI, Zugaro M (2022) Flexible communication between cell assemblies and ‘reader’ neurons. bioRxiv 2022.09.06.506754

6. Oberto VJ, Boucly CJ, Gao HY, Todorova R, Zugaro MB, Wiener SI (2022) Distributed cell assemblies spanning prefrontal cortex and striatum. Curr Biol 32:1–13. 10.1016/j.cub.2021.10.007

7. Russo E, Durstewitz D (2017) Cell assemblies at multiple time scales with arbitrary lag constellations. Elife 6:1–31. 10.7554/eLife.19428

8. Yang W, Yuste R (2017) In vivo imaging of neural activity. Nat Methods 14:349–359. 10.1038/nmeth.4230

9. Carrillo-Reid L, Yuste R (2020) Playing the piano with the cortex: role of neuronal ensembles and pattern completion in perception and behavior. Curr Opin Neurobiol 64:89–95. 10.1016/j.conb.2020.03.014

10. Carrillo-Reid L, Yang W, Bando Y, Peterka DS, Yuste R (2016) Imprinting Cortical Ensembles. Science (80-) 353:691–694

11. Chen TW, Wardill TJ, Sun Y, Pulver SR, Renninger SL, Baohan A, Schreiter ER, Kerr RA, Orger MB, Jayaraman V, Looger LL, Svoboda K, Kim DS (2013) Ultrasensitive fluorescent proteins for imaging neuronal activity. Nature 499:295–300. 10.1038/nature12354

12. Ali F, Kwan AC (2019) Interpreting in vivo calcium signals from neuronal cell bodies, axons, and dendrites: a review. Neurophotonics 7:1. 10.1117/1.nph.7.1.011402

13. Huang L, Ledochowitsch P, Knoblich U, Lecoq J, Murphy GJ, Reid RC, de Vries SEJ, Koch C, Zeng H, Buice MA, Waters J, Li L (2021) Relationship between simultaneously recorded spiking activity and fluorescence signal in gcamp6 transgenic mice. Elife 10:1–19. 10.7554/eLife.51675

14. Mölter J, Avitan L, Goodhill GJ (2018) Detecting neural assemblies in calcium imaging data. BMC Biol 16:1–20. 10.1186/s12915-018-0606-4

15. Gonzalez WG, Zhang H, Harutyunyan A, Lois C (2019) Persistence of neuronal representations through time and damage in the hippocampus. Science (80-) 365:821–825. 10.1126/science.aav9199

16. Carrillo-Reid L, Yang W, Kang Miller JE, Peterka DS, Yuste R (2017) Imaging and Optically Manipulating Neuronal Ensembles. Annu Rev Biophys 46:271–293. 10.1146/annurev-biophys-070816-033647

17. Wilson MA, McNaughton BL (1994) Reactivation of hippocampal ensemble memories during sleep. Science (80-) 265:676–679

18. Kudrimoti HS, Barnes CA, McNaughton BL (1999) Reactivation of hippocampal cell assemblies: Effects of behavioral state, experience, and EEG dynamics. J Neurosci 19:4090–4101. 10.1523/jneurosci.19-10-04090.1999

19. Humphries MD (2011) Spike-train communities: finding groups of similar spike trains. J Neurosci 31:2321–2336

20. Billeh YN, Schaub MT, Anastassiou CA, Barahona M, Koch C (2014) Revealing cell assemblies at multiple levels of granularity. J Neurosci Methods 236:92–106. 10.1016/j.jneumeth.2014.08.011

21. Peyrache A, Benchenane K, Khamassi M, Wiener SI, Battaglia FP (2010) Principal component analysis of ensemble recordings reveals cell assemblies at high temporal resolution. J Comput Neurosci 29:309–325

22. Lopes-dos-Santos V, Ribeiro S, Tort ABL (2013) Detecting cell assemblies in large neuronal populations. J Neurosci Methods 220:149–166. 10.1016/j.jneumeth.2013.04.010

23. Maboudi K, Ackermann E, de Jong LW, Pfeiffer BE, Foster D, Diba K, Kemere C (2018) Uncovering temporal structure in hippocampal output patterns. Elife 7:e34467

24. Mackevicius EL, Bahle AH, Williams AH, Gu S, Denisenko NI, Goldman MS, Fee MS (2019) Unsupervised discovery of temporal sequences in high-dimensional datasets, with applications to neuroscience. Elife 8:e38471

25. Cunningham JP, Yu BM (2014) Dimensionality reduction for large-scale neural recordings. Nat Neurosci 17:1500–1509. 10.1038/nn.3776

26. Freedman DA (2009) Statistical models: theory and practice. cambridge university press

27. Demars F, Todorova R, Makdah G, Forestier A, Krebs MO, Godsil BP, Jay TM, Wiener SI, Pompili MN (2022) Post-trauma behavioral phenotype predicts the degree of vulnerability to fear relapse after extinction in male rats. Curr Biol 32:3180–3188.e4. 10.1016/j.cub.2022.05.050

28. Pearson K. (1901) On lines and planes of closest fit to systems of points in space. Philos Mag 2:559–572

29. Jolliffe IT (2002) Principal component analysis for special types of data. Springer

30. Richmond BJ, Optican LM, Podell M, Spitzer H (1987) Temporal encoding of two-dimensional patterns by single units in primate inferior temporal cortex. I. Response characteristics. J Neurophysiol 57:132–146

31. Richmond BJ, Optican LM, Spitzer H (1990) Temporal encoding of two-dimensional patterns by single units in primate primary visual cortex. I. Stimulus-response relations. J Neurophysiol 64:351–369

32. Mcclurkin JW, Optican LM, Richmond BJ, Gawne TJ (1991) Concurrent Processing and Complexity of Temporally Encoded Neuronal Messages in Visual Perception. Science (80-) 253:675–677

33. Kjaer TW, Hertz JA, Richmond BJ (1994) Decoding cortical neuronal signals: Network models, information estimation and spatial tuning. J Comput Neurosci 1:109–139. 10.1007/BF00962721

34. Nicolelis MAL, Baccala LA, Lin RCS, Chapin JK (1995) Sensorimotor encoding by synchronous neural ensemble activity at multiple levels of the somatosensory system. Science (80-) 268:1353–1358. 10.1126/science.7761855

35. Chapin JK, Nicolelis MAL (1999) Principal component analysis of neuronal ensemble activity reveals multidimensional somatosensory representations. J Neurosci Methods 94:121–140. 10.1016/S0165-0270(99)00130-2

36. Marchenko VA, Pastur LA (1967) Distribution of eigenvalues for some sets of random matrices. Mat Sb 114:507–536

37. Peyrache A, Khamassi M, Benchenane K, Wiener SI, Battaglia FP (2009) Replay of rule-learning related neural patterns in the prefrontal cortex during sleep. Nat Neurosci 12:919–926. 10.1038/nn.2337

38. Benchenane K, Peyrache A, Khamassi M, Tierney PL, Gioanni Y, Battaglia FP, Wiener SI (2010) Coherent Theta Oscillations and Reorganization of Spike Timing in the Hippocampal-Prefrontal Network upon Learning. Neuron 66:921–936. 10.1016/j.neuron.2010.05.013

39. Gulati T, Ramanathan DS, Wong CC, Ganguly K (2014) Reactivation of emergent task-related ensembles during slow-wave sleep after neuroprosthetic learning. Nat Neurosci 17:1107–1113. 10.1038/nn.3759

40. Ramanathan DS, Gulati T, Ganguly K (2015) Sleep-Dependent Reactivation of Ensembles in Motor Cortex Promotes Skill Consolidation. PLOS Biol 13:e1002263. 10.1371/journal.pbio.1002263

41. Gulati T, Won SJ, Ramanathan DS, Wong CC, Bodepudi A, Swanson RA, Ganguly K (2015) Robust neuroprosthetic control from the stroke perilesional cortex. J Neurosci 35:8653–8661. 10.1523/JNEUROSCI.5007-14.2015

42. Tang W, Shin JD, Frank LM, Jadhav SP (2017) Hippocampal-prefrontal reactivation during learning is stronger in awake compared with sleep states. J Neurosci 37:11789–11805. 10.1523/JNEUROSCI.2291-17.2017

43. Rothschild G, Eban E, Frank LM (2017) A cortical–hippocampal–cortical loop of information processing during memory consolidation. Nat Neurosci 20:251–259. 10.1038/nn.4457

44. Chenani A, Sabariego M, Schlesiger MI, Leutgeb JK, Leutgeb S, Leibold C (2019) Hippocampal CA1 replay becomes less prominent but more rigid without inputs from medial entorhinal cortex. Nat Commun 10:1–13. 10.1038/s41467-019-09280-0

45. Sjulson L, Peyrache A, Cumpelik A, Cassataro D, Buzsáki G (2018) Cocaine Place Conditioning Strengthens Location-Specific Hippocampal Coupling to the Nucleus Accumbens. Neuron 98:926--934.e5. 10.1016/j.neuron.2018.04.015

46. Deolindo CS, Kunicki ACB, Brasil FL, Moioli RC (2014) Limitations of principal component analysis as a method to detect neuronal assemblies. In: 2014 IEEE 16th International Conference on e-Health Networking, Applications and Services (Healthcom). IEEE, pp 24–30

47. Brown GD, Yamada S, Sejnowski TJ (2001) Independent component analysis at the neural cocktail party. Trends Neurosci 24:54–63

48. Comon P (1994) Independent component analysis, a new concept? Signal Processing 36:287–314

49. Laubach M, Shuler M, Nicolelis MAL (1999) Independent component analyses for quantifying neuronal ensemble interactions. J Neurosci Methods 94:141–154

50. Giri B, Miyawaki H, Mizuseki K, Cheng S, Diba K (2019) Hippocampal reactivation extends for several hours following novel experience. J Neurosci 39:866–875. 10.1523/JNEUROSCI.1950-18.2018

51. Todorova R, Zugaro MB (2019) Isolated cortical computations during delta waves. Science (80-) 366:377–381

52. Trouche S, Koren V, Doig NM, Ellender TJ, El-Gaby M, Lopes-dos-Santos V, Reeve HM, Perestenko P V., Garas FN, Magill PJ, Sharott A, Dupret D (2019) A Hippocampus-Accumbens Tripartite Neuronal Motif Guides Appetitive Memory in Space. Cell 176:1393–1406.e16. 10.1016/j.cell.2018.12.037

53. Oliva A, Fernández-Ruiz A, Leroy F, Siegelbaum SA (2020) Hippocampal CA2 sharp-wave ripples reactivate and promote social memory. Nature 587:264–269. 10.1038/s41586-020-2758-y

54. Fernández-Ruiz A, Oliva A, Soula M, Rocha-Almeida F, Nagy GA, Martin-Vazquez G, Buzsáki G (2021) Gamma rhythm communication between entorhinal cortex and dentate gyrus neuronal assemblies. Science (80-) 372:eabf3119. 10.1126/science.abf3119

55. McKenzie S, Huszár R, English DF, Kim K, Christensen F, Yoon E, Buzsáki G (2021) Preexisting hippocampal network dynamics constrain optogenetically induced place fields. Neuron 109:1040–1054.e7. 10.1016/j.neuron.2021.01.011

56. El-Gaby M, Reeve HM, Lopes-dos-Santos V, Campo-Urriza N, Perestenko P V., Morley A, Strickland LAM, Lukács IP, Paulsen O, Dupret D (2021) An emergent neural coactivity code for dynamic memory. Nat Neurosci 24:694–704. 10.1038/s41593-021-00820-w

57. Almeida-Filho DG, Lopes-dos-Santos V, Vasconcelos NAP, Miranda JGV, Tort ABL, Ribeiro S (2014) An investigation of Hebbian phase sequences as assembly graphs. Front Neural Circuits 8:1–13. 10.3389/fncir.2014.00034

58. Bower MR, Stead M, Bower RS, Kucewicz MT, Sulc V, Cimbalnik J, Brinkmann BH, Vasoli VM, St Louis EK, Meyer FB, Marsh WR, Worrell GA (2015) Evidence for consolidation of neuronal assemblies after seizures in humans. J Neurosci 35:999–1010. 10.1523/JNEUROSCI.3019-14.2015

59. van de Ven GM, Trouche S, McNamara CG, Allen K, Dupret D (2016) Hippocampal Offline Reactivation Consolidates Recently Formed Cell Assembly Patterns during Sharp Wave-Ripples. Neuron 92:968–974. 10.1016/j.neuron.2016.10.020

60. Trouche S, Perestenko P V, Ven GM van de, Bratley CT, McNamara CG, Campo-Urriza N, Black SL, Reijmers LG, Dupret D (2016) Recoding a cocaine-place memory engram to a neutral engram in the hippocampus. Nat Neurosci 19:564–567. 10.1038/nn.4250

61. Conde-Ocazionez S, Altavini TS, Wunderle T, Schmidt KE (2018) Motion contrast in primary visual cortex: a direct comparison of single neuron and population encoding. Eur J Neurosci 47:358–369

62. Middleton SJ, Kneller EM, Chen S, Ogiwara I, Montal M, Yamakawa K, McHugh TJ (2018) Altered hippocampal replay is associated with memory impairment in mice heterozygous for the scn2a gene. Nat Neurosci 21:996–1003. 10.1038/s41593-018-0163-8

63. Deolindo CS, Kunicki ACB, da Silva MI, Lima Brasil F, Moioli RC (2018) Neuronal assemblies evidence distributed interactions within a tactile discrimination task in rats. Front Neural Circuits 11:114

64. See JZ, Atencio CA, Sohal VS, Schreiner CE (2018) Coordinated neuronal ensembles in primary auditory cortical columns. Elife 7:e35587

65. Guan H, Middleton SJ, Inoue T, McHugh TJ (2021) Lateralization of CA1 assemblies in the absence of CA3 input. Nat Commun 12:6114

66. Himberg J, Hyvärinen A, Esposito F (2004) Validating the independent components of neuroimaging time series via clustering and visualization. Neuroimage 22:1214–1222

67. Hyvärinen A, Ramkumar P (2013) Testing independent component patterns by inter-subject or inter-session consistency. Front Hum Neurosci 7:94

68. Klemm M, Haueisen J, Ivanova G (2009) Independent component analysis: comparison of algorithms for the investigation of surface electrical brain activity. Med Biol Eng Comput 47:413–423

69. Paatero P, Tapper U (1994) Positive matrix factorization: A non-negative factor model with optimal utilization of error estimates of data values. Environmetrics 5:111–126

70. Lee DD, Seung HS (1999) Learning the parts of objects by non-negative matrix factorization. Nature 401:788–791

71. Peter S, Kirschbaum E, Both M, Campbell L, Harvey B, Heins C, Durstewitz D, Diego F, Hamprecht FA (2017) Sparse convolutional coding for neuronal assembly detection. Adv Neural Inf Process Syst 30:

72. Tingley D, Buzsáki G (2020) Routing of Hippocampal Ripples to Subcortical Structures via the Lateral Septum. Neuron 105:138–149.e5. 10.1016/j.neuron.2019.10.012

73. Chen Z, Cichocki A (2005) Nonnegative matrix factorization with temporal smoothness and/or spatial decorrelation constraints. Lab Adv Brain Signal Process RIKEN, Tech Rep 68:

74. Choi S (2008) Algorithms for orthogonal nonnegative matrix factorization. Neural Networks 1828–1832

75. Hastie T, Tibshirani R, Friedman JH, Friedman JH (2009) The elements of statistical learning: data mining, inference, and prediction. Springer

76. Robotka H, Thomas L, Yu K, Wood W, Elie JE, Gahr M, Theunissen FE (2023) Sparse ensemble neural code for a complete vocal repertoire. Cell Rep 42:

77. Dejean C, Courtin J, Karalis N, Chaudun F, Wurtz H, Bienvenu TCM, Herry C (2016) Prefrontal neuronal assemblies temporally control fear behaviour. Nature 535:420–424. 10.1038/nature18630

78. Ghandour K, Ohkawa N, Fung CCA, Asai H, Saitoh Y, Takekawa T, Okubo-Suzuki R, Soya S, Nishizono H, Matsuo M, Osanai M, Sato M, Ohkura M, Nakai J, Hayashi Y, Sakurai T, Kitamura T, Fukai T, Inokuchi K (2019) Orchestrated ensemble activities constitute a hippocampal memory engram. Nat Commun 10:1–14. 10.1038/s41467-019-10683-2

79. Grosmark AD, Sparks FT, Davis MJ, Losonczy A (2021) Reactivation predicts the consolidation of unbiased long-term cognitive maps. Nat Neurosci 24:1574–1585

80. Pietri T, Romano SA, Pérez-Schuster V, Boulanger-Weill J, Candat V, Sumbre G (2017) The emergence of the spatial structure of tectal spontaneous activity is independent of visual inputs. Cell Rep 19:939–948

81. Girvan M, Newman MEJ (2002) Community structure in social and biological networks. Proc Natl Acad Sci U S A 99:7821–7826. 10.1073/pnas.122653799

82. Shimono M, Beggs JM (2015) Functional clusters, hubs, and communities in the cortical microconnectome. Cereb Cortex 25:3743–3757. 10.1093/cercor/bhu252

83. Pinotsis DA, Miller EK (2023) In vivo ephaptic coupling allows memory network formation. Cereb Cortex 33:9877–9895. 10.1093/cercor/bhad251

84. McMahon C, Kowalski DP, Krupka AJ, Lemay MA (2022) Single-cell and ensemble activity of lumbar intermediate and ventral horn interneurons in the spinal air-stepping cat. J Neurophysiol 127:99–115. 10.1152/jn.00202.2021

85. Arthur D, Vassilvitskii S (2007) K-means++: The advantages of careful seeding. Proc Annu ACM-SIAM Symp Discret Algorithms 07–09-Janu:1027–1035

86. Delvenne JC, Yaliraki SN, Barahon M (2010) Stability of graph communities across time scales. Proc Natl Acad Sci U S A 107:12755–12760. 10.1073/pnas.0903215107

87. Miyawaki H, Billeh YN, Diba K (2017) Low activity microstates during sleep. Sleep 40:. 10.1093/sleep/zsx066

88. Ponce-Alvarez A, Nácher V, Luna R, Riehle A, Romo R (2012) Dynamics of cortical neuronal ensembles transit from decision making to storage for later report. J Neurosci 32:11956–11969. 10.1523/JNEUROSCI.6176-11.2012

89. Mazzucato L, Fontanini A, La Camera G (2015) Dynamics of multistable states during ongoing and evoked cortical activity. J Neurosci 35:8214–8231. 10.1523/JNEUROSCI.4819-14.2015

90. Noguchi A, Huszár R, Morikawa S, Buzsáki G, Ikegaya Y (2022) Inhibition allocates spikes during hippocampal ripples. Nat Commun 13:. 10.1038/s41467-022-28890-9

91. Bollmann L, Baracskay P, Stella F, Csicsvari J (2023) Sleep stages antagonistically modulate reactivation drift

92. Oettl LL, Scheller M, Filosa C, Wieland S, Haag F, Loeb C, Durstewitz D, Shusterman R, Russo E, Kelsch W (2020) Phasic dopamine reinforces distinct striatal stimulus encoding in the olfactory tubercle driving dopaminergic reward prediction. Nat Commun 11:. 10.1038/s41467-020-17257-7

93. Pompili MN, Todorova R (2022) Discriminating Sleep From Freezing With Cortical Spindle Oscillations. Front Neural Circuits 15

94. Jun JJ, Steinmetz NA, Siegle JH, Denman DJ, Bauza M, Barbarits B, Lee AK, Anastassiou CA, Andrei A, Aydın Ç (2017) Fully integrated silicon probes for high-density recording of neural activity. Nature 551:232

95. Steinmetz NA, Aydin C, Lebedeva A, Okun M, Pachitariu M, Bauza M, Beau M, Bhagat J, Böhm C, Broux M, Chen S, Colonell J, Gardner RJ, Karsh B, Kloosterman F, Kostadinov D, Mora-Lopez C, O’Callaghan J, Park J, Putzeys J, Sauerbrei B, van Daal RJJ, Vollan AZ, Wang S, Welkenhuysen M, Ye Z, Dudman JT, Dutta B, Hantman AW, Harris KD, Lee AK, Moser EI, O’Keefe J, Renart A, Svoboda K, Häusser M, Haesler S, Carandini M, Harris TD (2021) Neuropixels 2.0: A miniaturized high-density probe for stable, long-term brain recordings. Science (80-) 372:. 10.1126/science.abf4588

96. Ghestem A, Pompili MN, Dipper-Wawra M, Quilichini PP, Bernard C, Ferraris M (2023) Long-term near-continuous recording with Neuropixels probes in healthy and epileptic rats. bioRxiv 1–19. 10.1101/2023.02.16.528689

97. Pandarinath C, O’Shea DJ, Collins J, Jozefowicz R, Stavisky SD, Kao JC, Trautmann EM, Kaufman MT, Ryu SI, Hochberg LR, Henderson JM, Shenoy K V., Abbott LF, Sussillo D (2018) Inferring single-trial neural population dynamics using sequential auto-encoders. Nat Methods 15:805–815. 10.1038/s41592-018-0109-9

